# Mutational oncogenic signatures on structurally resolved protein interacting interfaces

**DOI:** 10.1101/016204

**Authors:** Luz Garcia-Alonso, Joaquin Dopazo

**Author notes:** Present address: European Bioinformatics Institute (EMBL-EBI), European Molecular Biology Laboratory, Wellcome Trust Genome Campus, Hinxton, United Kingdom.

## Abstract

The importance of the context of interactions in the proteins mutated in cancer is long known. However, our knowledge on how mutations affecting to protein-protein interactions (PPIs) are related to cancer occurrence and progression is still poor. Here, we extracted the missense somatic mutations from 5920 cancer patients of 33 different cancer types, taken from the International Cancer Genome Consortium (ICGC) and The Cancer Genome Atlas (TCGA), and mapped them onto a structurally resolved interactome, which integrates three-dimensional atomic-level models of domain-domain interactions with experimentally determined PPIs, involving a total of 7580 unique interacting domains that participate in 13160 interactions connecting 4996 proteins. We observed that somatic nonsynonymous mutations tend to concentrate in ordered regions of the affected proteins and, within these, they have a clear preference for the interacting interfaces. Also, we have identified more than 250 interacting interfaces candidate to drive cancer. Examples demonstrate how mutations in the interacting interfaces are strongly associated with patient survival time, while similar mutations in other areas of the same proteins lack this association. Our results suggest that the perturbation caused by cancer mutations in protein interactions is an important factor in explaining the heterogeneity between cancer patients.

## Introduction

Aberrant protein activities caused by mutations altering protein-protein interactions have been associated to cancer initiation and progression. A well-known example is the Y42C mutation in BRCA2 that inhibits its interaction with the protein A, essential for DNA repair, replication and recombination, leading to the accumulation of DNA damage (Wong et al., 2003). From a global point of view, systematic analysis of the role cancer driver genes in PPI networks (Goh et al., 2007; Feldman et al., 2008) depicted a clear association of these to code for highly connected and central proteins, suggesting that the impairment of protein interactions at the core of the interactome can be a common mechanism in tumor evolution. This is in agreement with our previous observation that somatic mutations from CLL patients tend fall in central modules of the interactome (Garcia-Alonso et al., 2014), In fact, the products of cancer driver genes participate in a greater number of interactions (network hubs) (Jonsson and Bates, 2006) and are more likely to influence a larger number of biological processes (pleiotropy) (Yu et al., 2008).

These observations, although pointing at the global properties of genes involved in cancer, can only be seen as descriptive rather than propose testable hypotheses. The reason behind is the lack of molecular details in the way the interactome is modeled (ie. an undirected graph), where proteins represent graph-theoretical nodes ignoring its structural details. However, not all the mutations from the same protein have the same consequence. The impact of a mutation depends on the stereochemical nature of the change and, ultimately, on its location within the carrier protein, which evidences the importance of integrating the structural properties in approaches aimed to decipher the effect of cancer mutations on protein coding genes.

Here we investigate the relevance of protein interacting interfaces in the tumorigenic process. We hypothesize that somatic mutations are more likely to confer a functional change and, therefore, to be selected if they alter a molecular interaction, specially when occur in proteins that govern essential biological processes. For this purpose, we have structurally resolved the human interactome by integrating three-dimensional (3D) atomic-level models of domain-domain interactions with experimentally determined known, and mapped the somatic mutations from 5920 cancer patients onto it. First, we have performed a global pan-cancer analysis to study the distribution of nonsynonymous mutations among the different structural protein regions and the interactome location. Encouraged by the results, we next applied a systematic protein-centric strategy to detect specific protein interacting interfaces containing an unexpected amount of mutations with respect to the number of mutations in the protein sequence. For some detected interfaces we observe a clear relationship with the patient survival time, highlighting the relevance of considering the mutation location when studying cancer genetic variation. Finally, we mapped the enriched PPI interfaces onto the 3D interactome to provide a high-resolution picture of cancer mutations that describes well-known tumorigenic processes but also proposes novel mechanistic hypotheses in cancer.

## Results and discussion

### Construction of a 3D structurally resolved protein interactome

The starting point of this study is to resolve the 3D structure of the PPIs from the interactome. Here, we followed the homology modeling approach proposed by Wang and colleges (Wang et al., 2012). Specifically, we considered all binary interactions from our curated interactome used in our previous study (Garcia-Alonso et al., 2014) that contain a Pfam domain pair interacting in at least one co-crystal structure in the PDB and, therefore, the specific interactor interface of each participant is known. That is, we use co-crystal structures as a gold-standard proof that these interactions do occur. The obtained 3D interactome consisted of a total of 7580 unique interacting domains in 13160 PPI between 4996 proteins. This new version of the interactome provides a high-resolution and accurate description of the molecular interactions and is a valuable resource for interpreting the massive amount of genomic data generated from thousands of patients.

### Relevance of the protein interacting interfaces in cancer

Next, we explored the distribution of the somatic mutation among the protein interacting Pfams. In general, Pfam domains have a greater overlap with ordered regions (Bateman et al., 2002). Since the disordered/unstructured regions of proteins are less restricted spatially and, without some exceptions (Chen et al., 2006), evolve more rapidly than ordered/structured regions (Brown et al., 2002), we expect that more somatic mutations would occur in these regions. This unequal evolutionary constraint, observable in the proteins of the interactome (*P* ≤ 0.001, Wilcoxon test, Figure 1A), can bias the results in such a way that the enrichment of mutations in the interacting Pfam domains would be underestimated. To overcome this bias, we calculated the probability estimate of each residue in the sequence being disordered and split the protein sequence in ordered/disordered regions (Figure 1B).

**Figure 1.**
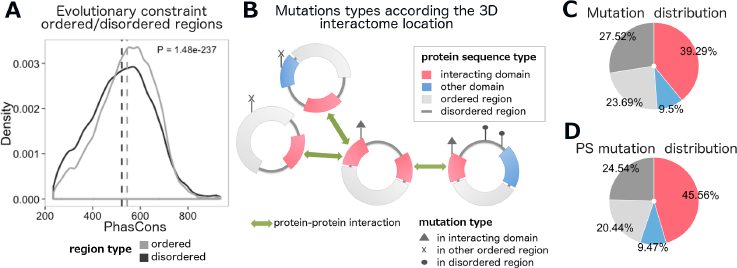
Mapping of cancer nonsynonymous mutations in the structurally resolved protein interfaces. **A** Evolutionary constraint distribution for somatic nonsynonymous mutations in ordered (light grey) and disordered (dark grey) protein sequences. Significance p-value for the comparison is displayed on the top corner (Wilcoxon test). **B** Classification of cancer nonsynonymous mutations according its location in the structurally resolved proteins of the interactome. **C** and **D** Proportions of, respectively, all and positively selected (PS) cancer nonsynonymous mutations among interacting Pfam domains, non-interacting Pfam domains, other ordered and disordered protein sequences.

After merging the somatic exonic data and filtering donors with no somatic variants in the proteins of the interactome, we studied the exome sequence of 5920 cancer donors from 33 different cancer types. We found a total of 176316 nonsynonymous somatic mutations, mapping 4846 proteins in the interactome. 69563 (39.29%) of these mutations were located within the interacting Pfam interfaces, 16822 (9.5%) in other Pfam domains, 41949 (23.69%) in other ordered regions and 48732 (27.52%) in disordered regions (Figure 1C). These mutations were tested for patterns of positive selection using both oncoDriveFM and oncoDriveCLUST methods, obtaining a total of 22543 variants under positive selection in the tumorigenic process: 10270 (45.56%) in interacting Pfams, 2134 (9.47%) in other Pfams, 4607 (20.44%) in other ordered regions and 5532 (24.54%) in disordered regions (Figure 1D).

Next, we aimed to investigate whether cancer somatic mutations were differently distributed among the distinct protein regions. First, we studied the preferential location of nonsynonymous mutations for either ordered or disordered regions by comparing the observed number of mutations in the ordered regions for all interactome proteins to the expected distribution, obtained by permuting the variants among the whole protein sequences (Figure 2A). Results show enrichment for the somatic mutations in ordered regions (*P* ≤ 0.001, Permutation test). Second, we focused on the distribution among interacting domains. Here we also detect a significant overrepresentation of cancer mutations (*P* ≤ 1.001, Permutation test), compared against the random expectations given by the mutation frequency among ordered regions (Figure 2B). When we divided the mutations according whether they were predicted to be under positive selection or not, we observe the same patter for both group of mutations (*P* ≤ 0.001 both, Permutation test). Finally, we focused on the when looking at the noninteracting domains (Figure 2C). Somatic mutations are significantly underrepresented at these regions (*P* ≤ 0.001, Permutation test) whereas no significance was found for mutations under positive selection (*P* = 0.16, Permutation test).

**Figure 2.**
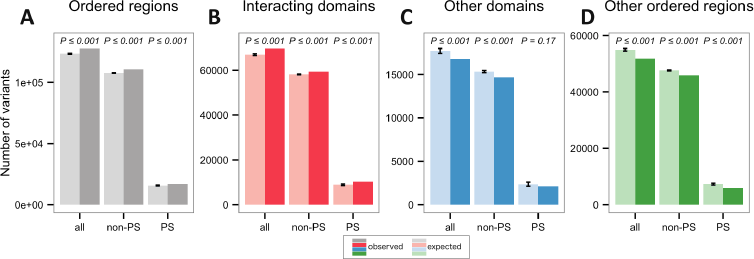
Distribution of cancer nonsynonymous mutations among the structurally resolved protein region types. The bars represent the total number of nonsynonymous mutations observed (dark color) and expected (light color) in ordered protein sequences (**A**), interacting Pfam domains (**B**), non-interacting Pfam domains (**C**) and other ordered regions (**D**). P-values are shown on top of bars (Permutation test). For the study of the mutation frequency in the ordered protein sequences (A), the distribution of the expected values was obtained by permuting mutations across the whole protein sequence (considering both ordered and disordered regions). For the study of the mutation frequency in the PPI interfaces, non-interacting domains and other ordered regions (B, C and D), the distribution of the expected values was obtained by permuting mutations across the ordered protein sequence. Expected error bars represent standard errors of the mean value from permutations.

Focusing on the interacting interfaces, we next classified the interacting interfaces according to whether they occur (i) in proteins that occupy the periphery of the interactome (defined as the proteins that fall in the first quartile of the distribution for the closeness centrality parameter, Q1), (ii) in proteins with an intermediate centrality (in the second and third quartiles, Q2-Q3) or (iii) central proteins (in the fourth quartile, Q4) Then the same test performed in the Figure 2B was repeated for each protein group. The results display a more precise and opposite signal between the mutations under positive selection and the rest of mutations (Figure 3). The mutations under positive selection significantly concentrate in the interacting interfaces of the central proteins (*P* ≤ 0.001, Permutation test), confirming that the impairment of the central binding interactions is a positively selected network hallmark in cancer development. In contrast, the rest of the mutations display a tendency toward the network periphery, behaving similarly to the distribution of the germinal variants from healthy individuals (Garcia-Alonso et al., 2014).

**Figure 3.**
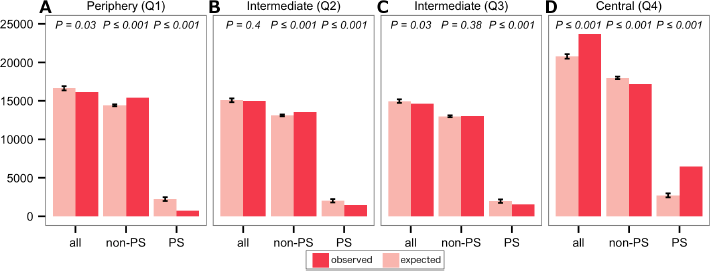
Distribution of nonsynonymous mutations among structurally resolved PPI interfaces according to the network centrality. The bars represent the total number of nonsynonymous mutations observed (dark color) and expected (light color) in PPI interfaces for proteins located in the periphery of the interactome (defined as the proteins that fall in the first quartile of the distribution of the closeness centrality parameter, Q1), in intermediate regions (in the second and third quartiles, Q2-Q3) or occupying central positions (in the fourth quartile, Q4). P-values are shown on top of bars (Permutation test). The distribution of the expected values was obtained by permuting mutations across the ordered sequences of the interactome proteins. Expected error bars represent standard errors of the mean value from permutations.

### Identifying significantly mutated protein interacting interfaces

Motivated by the enrichment of cancer mutations in the binding interfaces of the interactome and its potential functional implications for cancer development, we aimed to search for proteins with a bias in their mutation rates towards its interacting interfaces, which we hypothesize would contribute to tumor evolution. Thus, focusing on each specific interacting domain, we performed a protein-centric mutation enrichment analysis with one tailed binomial test. We consider that FDR P-values ≤ 0.05 indicate that the ratio of observed mutations in the interacting domain is significantly greater than the ratio of mutation along the ordered protein sequence. Interfaces with less than 5 mutations were discarded. Additionally, as the mutation rates are not homogeneous across the genome (Liu et al., 2013) and our results could be biased towards hyper-mutated sequence regions, we performed a second test over the synonymous mutations, which are not expected to have a functional implication at the protein level. We assume that those interacting interfaces that are also enriched in synonymous mutations follow the baseline distribution of somatic mutations and, therefore, were discarded since we cannot attribute a direct functional implication.

Systematic analysis per cancer type reveals 83 (FDR *P* ≤ 0.05) proteins concentrating its somatic mutations on the interacting interfaces (Figure 4). Further analysis across the merged pan-cancer dataset leads to the identification of 161 additional proteins (Figure 5), which sums up a total of 252 significantly enriched interacting interfaces in 248 proteins (see supplementary table 1). These predicted interfaces encode potential molecular mechanism for a total of 4308 nonsynonymous mutations.

**Figure 4.**
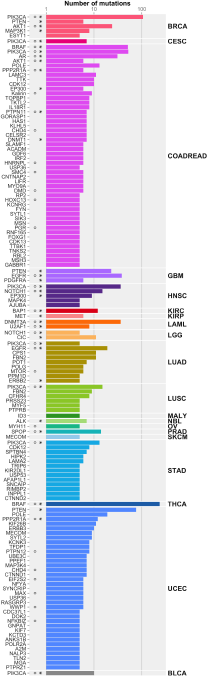
Proteins with significantly mutated interacting interfaces per cancer type. Barplot shows the number of nonsynonymous mutations for each enriched Pfam-protein colored based on the cancer type. Proteins are ordered according to the decreasing frequency of mutations in the interacting interface.

**Figure 5.**
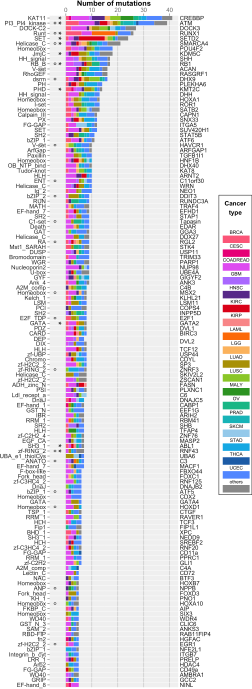
Significantly mutated protein interacting interfaces in pancancer. Barplot shows the number of nonsynonymous mutations for each enriched Pfam-protein colored based on the cancer type. Proteins are ordered according to the decreasing frequency of mutations in the interacting interface.

#### Most known cancer genes accumulate mutations on the interacting interfaces

Among the identified proteins, 32 (13%) are in the Cancer Driver list proposed by Vogelstein et al. (2013) (*P* = 4.25 x 10^-18^, Fisher’s exact test; genes labelled with * in Figure 5). Also, over 38.80% of the mutations in the identified interacting interfaces are considered under positive selection by the oncoDriveFM and oncoDriveCLUST predictors (*P* ≤ 0.001, Fisher’s exact test), whereas the rest 61.19% of mutations would imply new mechanisms.

Functional profiling of the enriched proteins using identifiedO tool (Al-Shahrour et al., 2004) from Babelomics (Medina et al., 2010) reveals an overrepresentation (compared to the whole interactome members) of processes and pathways that are hallmarks of cancer (Hanahan, 2000; Hanahan and Weinberg, 2011), such as regulation of gene expression, regulation of the cell cycle and apoptosis, chromatin modification, protein processing, tyrosine kinase signalling pathways and KEGG pathways in cancer (FDR *P* ≤ 0.05, Fisher’s exact test). This observation demonstrates that the presented strategy is useful to propose new candidate drivers.

Focusing on the individual predicted interfaces, our results highlight, for example, the pleckstrin homology (PH) kinase domain in AKT1 (Figure 6A), a member of the AKT protein complex, that links several key processes including metabolism, proliferation, cell survival, growth and angiogenesis. PH domain-kinase domain interactions are necessary in maintaining AKT in an inactive state through autoinhibitory interactions and mutations in the PH-kinase interface constitutively active AKT, whose aberrant activity leads to cellular transformation (Parikh et al., 2012). These AKT1 mutants are not effectively inhibited by allosteric AKT (currently under investigation in preclinical and clinical testing) evidencing that the mutational status of PH domain in AKT has important implications for the choice of treatment in the clinic (Calleja et al., 2009; Wu et al., 2010). Another example is the meprin and traf homology (MATH) domain in SPOP protein (Figure 6B), an ubiquitin ligase that promotes the ubiquitin-mediated degradation of the proteins binding to its substrate recognition domain. SPOP substrates are proteins implicated in transcriptional regulation of genes involved in essential cellular functions. Some examples among its substrates are NCOA2 and NCOA3, master activators of several transcription factors such as AR, a well-known cancer driver gene (Li et al., 2011; Geng et al., 2013).

**Figure 6.**
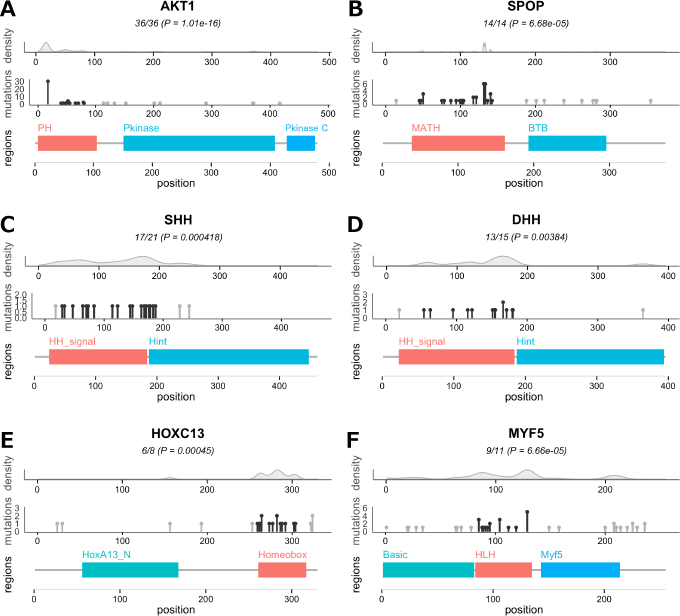
Examples of proteins with enriched interacting domain. For each protein, three plots are displayed. From top to bottom: (1) mutation density among ordered protein sequence, (2) total mutation counts for the predicted PPI interface (black) and other protein regions (gray); and (3) location of Pfam domains among the protein sequence. The predicted interacting interface is colored in red.

#### Novel candidate genes accumulating mutations on the interacting interfaces

Our test highlights novel candidates such as SHH and DHH, members of the Hedgehog (Hh) pathway, or MYF5 and HOXC13, both transcription factors (Figure 6C-F). SHH and DHH proteins are of special interest since they are critical regulators for tissue differentiation and, in adulthood, are involved in the maintenance of homeostasis. Aberrant Hh pathway activity has been implicated in a broad variety of tumors and has been hypothesised to play an important role in the formation and maintenance of cancer stem cells (CSCs). Although it has become clear that aberrant activity of Hh pathway either by point mutations of the downstream proteins (PTCH1, SMO or SUFU) or ligand over-expression leads oncogenic signalling (Ruch and Kim, 2013), to our knowledge, few mutations affecting directly the sequence of the SHH and DHH have been proposed as a tumorigenic mechanism (Oro et al., 1997). The cause may be the relatively low frequency of its mutations when each cancer type is studied separately. This highlights that integrated pan-cancer approach can be crucial in the detection of new cancer mechanisms. Particularly, the altered domain is responsible for the HH signal and directly binds to HHIP, which regulates Hh signalling negatively (Chuang and McMahon, 1999). Therefore we propose that the impairment of the binding of SHH and DHH with its inhibitor could be a hypothetical mechanism for cancer, although further studies should be conducted to corroborate its implications in cancer.

#### Approach comparison

Opposite to the conventional methods based on mutation frequency that search for highly recurrently mutated genes, our strategy studies each gene separately. As observed in other ‘‘genecentric” (Gonzalez-Perez and Lopez-Bigas, 2012; Tamborero et al., 2013; Reimand and Bader, 2013), our approach is able to detect individual interacting interfaces whose mutational rate is low but unexpected given the protein-wide number of mutations. In fact, our results include products from the long tail of genes with low frequency mutations (ex. SHH and DHH, members of the Hedgehog (Hh) pathway, or MYF5 and HOXC13, both transcription factors). After this approach was designed and applied to the cancer data, we learned about the publication of a similar study by Porta-Pardo and Godzik (2014) called e-Drive. Similarly as the approach we propose here, e-Drive studies the accumulation of cancer mutations on pre-defined gene functional regions using one tailed binomial tests, which supports the use of this test for such objective. However, e-Drive does not correct for the accumulation of synonymous variants and, therefore, hyper-mutated interfaces that do not confer selective advantage are not distinguished.

Yet, our approach has a major limitation: genes for which mutations are homogeneously distributed across the sequence, such as tumor suppressors, cannot be detected with our method since it specifically search for deviations from even distribution. For example, loss-of-function mutations in the key tumor suppressor gene P53 are scattered across its sequence and, although is the gene with the highest number of mutations, our method does not detect it. Thus, our method is complementary to the overall frequency based methods and the capture of all the players in oncogenesis requires the combination of the different strategies.

### Clinical relevance of mutations affecting interacting interfaces: survival implications

To explore the prognosis significance of the oncogenic candidate interacting interfaces, we downloaded survival clinical data from Breast Cancer (BRCA) patients. Breast cancer is the most commonly diagnosed cancer among women in Europe, causing 131,200 deaths in 2012 (Ferlay et al., 2013). Our aim at this step is to study the role of the mutations located on the interacting interfaces with the tumor evolution. Our hypothesis is that if hampering an interaction is a relevant mechanism for the tumor progression, then mutations located at the interface (and mutations that considerably change the overall protein structure) would display a significant impact on survival in the carriers.

For each interacting interface, we split the patients into two groups: one containing mutations in the interacting interface and the other with mutations in the same protein but outside the interacting interface. Results show that mutations in the PI3Ka domain (exon 9) of PIK3CA were strongly associated with increased survival compared to the rest of mutations (Figure 7A). However, a bibliography research reveals that the prognostic value of PIK3CA mutations at different regions in breast cancer remains controversial. Whereas some studies have reported that the presence of H1047R (PI3 PI4 kinase, exon 20) mutation was strongly associated with the absence of lymph node metastasis and better prognosis (Barbareschi et al., 2007; Kalinsky et al., 2009), our results and three other studies report that exon 9 mutations are associated with increased survival in breast cancer (Lai et al., 2008; Mangone et al., 2012; Arsenic et al., 2014) and lung adenocarcinoma (Zhang et al., 2013). Splitting patients into fourth groups (with mutations in the predicted PPI interface, H1047R mutation, mutations in the rest of PIK3CA and no mutations in PIK3CA) reveals a difference in survival between donors (Figure 7B-C), being the patients with mutation in the predicted interacting interface those with better prognosis. The difference is maintain when the information of the direct interactors is included, although due to the low recurrence of the latest, an independent validation of the mutations in PI3KR1 and PI3KR3 can not be assessed (Figure 7D). These mutations were previously investigated in Glioblastoma in primary studies from the TCGA, where it was proposed that mutations on the PIK3a domain (exon 9) might prevent inhibitory contact with PI3KR1 and PI3KR3, causing constitutive PI3K activity (McLendon et al., 2008) (Figure 7E).

**Figure 7.**
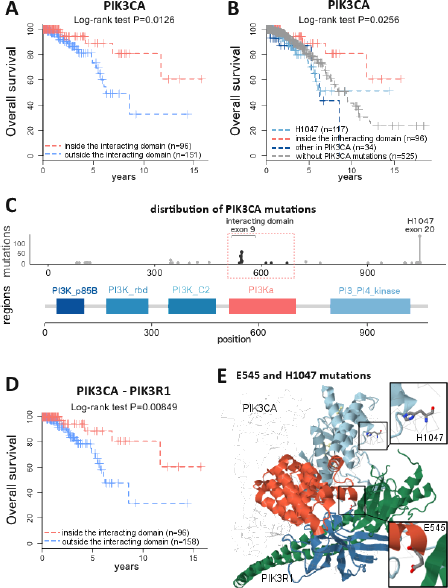
Survival analysis on PIK3CA mutations in BRCA patients. **(A)** Kaplan-Meier survival plots for patient with mutations inside (red) or outside (blue) the PIK3CA interacting domain. **(B)** Kaplan-Meier survival plots for patient with mutations inside the PIK3CA interacting domain (red), H1047 mutation (light blue), other PIK3CA mutations (dark blue) or wild type PIK3CA (gray). **(C)** Location of mutations among PIK3CA sequence domains **(D)** Kaplan-Meier survival plots for patient with mutations inside (red) or outside (blue) the PIK3CA-PIK3R1 interacting domains **(E)** Mutations location in the 3D structure of the interaction between PIK3CA (domains in orange and light blue) and PIK3R1 (domains in green and dark blue). Model extracted from PDB ref. 4JPS

Several clinical variables can account for the discrepancies among studies in the role of mutations from different exons, such as the tumor subtype, stage, surgery, therapeutic treatment or even population heterogeneity. Additional studies on larger and more homogeneous series of patients are necessary to verify the real significance of the association. Nevertheless, results evidence that mutations in different PIK3CA regions may play different roles in tumor evolution.

### Topological properties of enriched interfaces

Our work would be incomplete without an analysis of the topological properties of the affected interfaces. The basic questions to assess are: Do the affected interfaces have a preference to occupy central positions? Do they involve more interactions than the non-affected interfaces? Are they more likely to interact with one another, or are they spread around the interactome? To address these questions, we mapped the enriched domains onto our structurally resolved protein interactome and examined its topological properties. As expected, 33.66% of the proteins with predicted PPI interfaces are located in central positions (Q4, *P* = 0.0138, Fisher’s exact test). Next, interested in deciphering if the predicted interfaces were concentrating more interactions, we observed that enriched interfaces participate in more interactions than the interfaces of non-enriched proteins but (*P* = 0.0333, Wilcoxon test). Interestingly, no difference was found compared to non-enriched interacting interfaces of the same predicted proteins (*P* = 0.479, Wilcoxon test), which suggests that the hub role of cancer genes seems a property relative to the whole protein more than to the enriched interface (Figure 8B).

Biological processes involved in oncogenesis converge in common regulators that modulate crosstalk between them (Bustin, 1998). To study whether this property is reproduced in our predicted interfaces, we look for predicted interfaces that interact directly and observed a total of 37 interactions in which both interfaces in the PPI are enriched in mutations. To evaluate if the predicted interfaces have a higher tendency to interact with one another, we permuted 1000 times the interactor labels of the network while preserving the total connectivity of each protein. Results show that in only 0.3% of the random cases we obtain a value equal or greater than the observed (*P* = 0.003, Permutation test, Figure 8C). This analysis was repeated using the non-enriched interfaces of the same predicted proteins and observed 25 direct interactions, and no significant difference from the random expectation was detected (*P* = 0.73, Permutation test, Figure 8D).

Motivated by the results, we remodeled the interactome into an undirected graph by an alternative approach in which nodes represent protein interfaces (instead o the whole protein) and edges the interactions either between or within proteins (Figure 8E). This new version of the interactome incorporates a new level of complexity that discriminates between different edges from the same protein when interact through different interfaces. In order to study the crosstalk between enriched and non-enriched PPI interfaces, all occurring in the predicted proteins, we computed the distances within each set of interfaces by means of the average length of the shortest path. We found that the distribution of shortest network distances is skewed towards shorter paths for the enriched interfaces, which can be a consequence of this preferential interaction affinity of the enriched interfaces to interact with each other as compared with non-enriched (Figure 8F). The relevance of this observation is that it points out at the function-centric organization of the interactome, where different proteins can cause similar clinical disorders when they affect regions regulating common functions.

**Figure 8.**
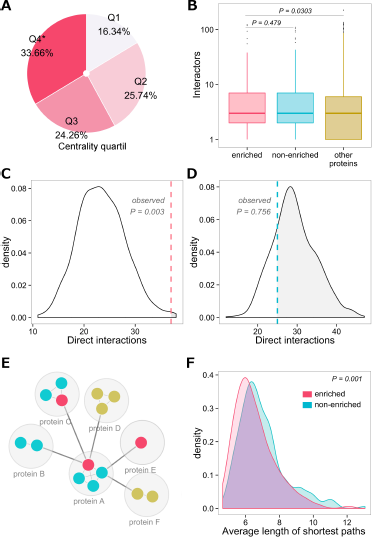
Topological properties of affected interfaces. **(A)** Proportion of proteins with enriched interfaces in each centrality quartile. **(B)** Interaction degree comparison between enriched interfaces, non-enriched interfaces in the same predicted proteins and interfaces in non-predicted proteins. **(C)** Expected distribution and observed value (dashed red line) for the number of direct partners between the enriched interfaces. **(D)** Expected distribution and observed value (dashed blue line) for the number of direct partners between the non-enriched interfaces in the same predicted proteins. **(E)** Graph model of the structurally resolved protein interactome. **(F)** Average length distribution of the shortest paths within enriched interfaces (red) and non-enriched interfaces from the same proteins.

### The 3D cancer interactome: new insights into the cancer hallmarks

To extract a rationale map of the proteins with driver interacting interfaces, we calculated the MCN allowing one intermediate using the SNOW tool (Minguez et al., 2009). The identified subnetwork involves 535 interactions between 293 proteins (153 are external intermediates added to connect the network, and 15 of them are known-driver genes) (Figure 9). As commented before, the subnetwork contains 39 direct interactions between predicted interfaces of 37 proteins (MCN, *P* = 0.019, SNOW Permutation test), that is, interactions in which both PPI interfaces are enriched in somatic mutations (bold edges in Figure 9B). Over the 56.12% of the donors have at least one somatic mutation in the interfaces of the MCN PPIs.

**Figure 9.**
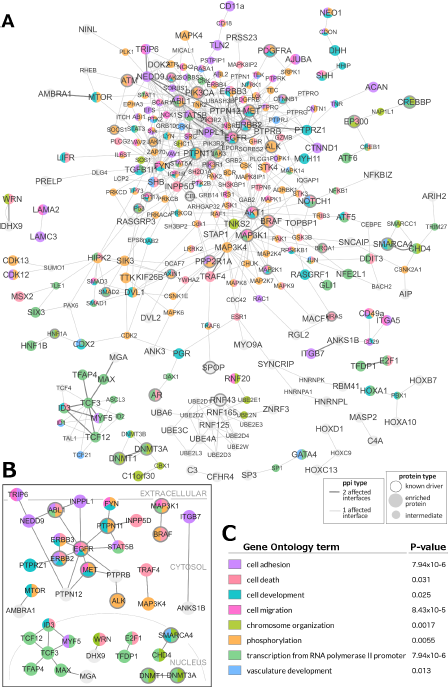
3D subnetwork of enriched interacting domains. **(A)** MCN between the enriched interfaces allowing one intermediate. **(B)** MCN between the enriched interfaces allowing without intermediates. **(C)** Main GO Biological Processes enriched in the MCN.

Several well-known cancer processes and pathways are embedded in the predicted subnetwork, such as the PIK3-AKT, the ErbB, Jak-STAT, AKT/mTOR signalling pathway, PTEN dependent cell cycle arrest and apoptosis, the DNA-dependent binding transcriptional regulators, etc, which validates our approach. Again, when performing a functional enrichment over both affected proteins and direct partners, we observe an overrepresentation (compared to the whole interactome members) of processes and pathways that are hallmarks of cancer, such as transcription factors, chromatin modificators, phosphorylation processes, regulators of cell development and death, and even proteins implicated in the vascular development and angiogenesis (FDR *P* ≤ 0.05, Fisher’s exact test).

However, this subnetwork provides also novel hypotheses for regulation of transcription processes potentially involved in tumor progression. One example is the network component formed by the members of basic helix-loop-helix (bHLH) family: ID3, MAX, MGA, TFAP4, TCF12, TCF3, MYF5 (Figure 10A-B). All these proteins are members of a well-known group of transcriptional regulators of genes involved in the regulation of gene expression and cell fate. They form homo/heterodimers by the non-covalent interaction between the bHLH domains, which is required for an efficient DNA binding. Interestingly, three recent exome sequencing studies of Burkitt’s lymphoma patients provided convincing support for the idea that ID genes may function as tumor-suppressors. Concretely, mutations affecting TCF3 or its negative regulator ID3 are found in the 70% of sporadic Burkitt’s lymphoma, blocking the interaction between TCF3 and ID3 breaking the negative regulatory loop created by ID3 (Figure 10C) (Schmitz et al., 2012; Richter et al., 2012). Moreover, other studies have described a crosstalk between the HH and WNT/*β*-catenin pathways, which cooperate inducing expression of some bHLH proteins in cancer (Javelaud et al., 2011). In this sense, HH signal activates bHLH family expression during development and inappropriate activation of bHLH signaling in individual cells may contribute to tumor initiation, as observed for Rhabdomyosarcoma (Gerber et al., 2006).

**Figure 10.**
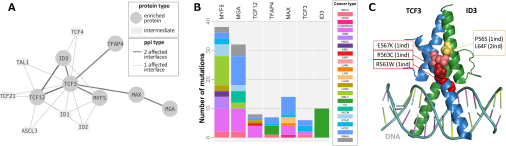
Cancer mutations in interacting domains in bHLH family of TFs. Cancer mutations in interacting domains in bHLH family of TFs. **(A)** Interacting interfaces of bHLH enriched in cancer mutations. Dark grey nodes indicate proteins carrying the enriched interfaces. Dark grey edges indicate interactions where both interfaces are enriched **(B)** Significantly mutated bHLH TFs interacting interfaces per cancer type. Barplot shows the number of mutations for each enriched Pfam-protein colored based on the cancer type. Proteins are ordered according to the decreasing frequency of mutations in the interacting interface. **(C)** Mutations from TCF3 and ID3 on the 3D structure of bHLH domain predicted to impact the dimer interacting interface (Model extracted from PDB ref. 2LFH)

As for the HH pathway members, much of the previous work has focused on studying the gene expression patterns of the bHLH family in cancer but, to our knowledge, mutations in the subnetwork formed by the bHLH TFs have not been directly implicated yet. As described before, only few works have found an association between ID3, TCF12 and TCF3 proteins and tumor initiation. Our results suggest that mutations in this network component may cause aberrant expression of genes involved in proliferation/cell fate determination by affecting binding events between these TFs and, thus, contribute to the progression of the malignant phenotype (Figure 10C).

## Conclusions

First, our exploration of the distribution of cancer somatic mutations among the regions of the proteome showed a clear pattern: the mutations tend to concentrate in ordered regions of the proteins and, within these, they have a clear preference for the interacting interfaces. Moreover, when the centrality of the protein within the interactome is considered, we observed a clear tendency of the somatic mutations to occur at proteins that occupy central positions. Finally, when we divide the mutations into two groups, one including the subset of mutations under positive selection and other including the rest of mutations, we observe a more specific and opposite pattern between them. Mutations under positive selection concentrate in central regions and avoid the periphery, while the rest of mutations concentrate at the interactome periphery. Thus, cancer mutations under positive selection follow the opposite pattern to that observed in population deleterious variants of apparently healthy individuals (Garcia-Alonso et al., 2014).

Also we propose a new approach to identify somatic mutations whose effect may be advantageous for cancer development. By studying the mutation distribution among the functional regions within each protein, we have identified more than 250 interacting interfaces candidate to drive cancer. Several sources of evidence support the potential role of the identified mutations: first, they accumulate in a specific region instead of being homogeneously distributed among the protein; second, these regions are conserved, which functions have been evolutionary maintained; and third, these regions mediate protein-protein interactions, which are crucial in mediating the transmission of the biological signals. Thus, these interacting interfaces can be seen as mechanistic hypotheses candidates to explain the molecular basis behind the genetic-cancer associations for most of the known cancer genes. This fact is the most significant difference from frequency-based methods, which does not add functional insight into the identified genes.

The power of the approach is evidenced by the ability of detecting low frequency mutations, which direct role in oncogenesis is sustained by independent published studies. Examples of these mutations are those falling in the HH domain of SHH and DHH (Oro et al., 1997), in the homeobox domain in HOXC13 and in the bHLH domain of the bHLH family of transcription factors (Schmitz et al., 2012; Richter et al., 2012). Moreover, the approach identified interacting interfaces which mutational state relates to patients prognosis, being able to explain survival differences in patients of the same 16 tumor type and with the same mutated gene (Lai et al., 2008; Mangone et al., 2012; Arsenic et al., 2014).

Taken all together, our results suggest that qualitative changes in protein interactions could explain heterogeneity between cancer patients better than considering isolated genes, ignoring the structural details that mediate its communication with other molecules in the context of the interaction network. In a scenario where is becoming evident the heterogeneity between cancer patients and the complexity of predicting prognosis outcome, zooming into the molecular consequences of genomic variants and contextualizing them in the network of molecular interactions would constitute a step forward towards personalized medicine.

## Materials and Methods

### Construction of the structurally resolved human protein interactome

Here we aimed to structurally resolve the protein interactome used in the a previous study (Garcia-Alonso et al., 2014) by predicting the protein interaction interfaces of each ppi. For this purpose, we followed the homology modeling approach proposed by Wang and colleges (Wang et al., 2012) where cocrystal structures are used as a gold-standard evidence to resolve the structural details of PPIs. For each protein from the interactome, we used the protein domain definition proposed in Pfam (Finn et al., 2013) and retrieved the protein sequence-Pfam mappings from the Uniprot database (Consortium, 2011). Next, we collected from 3did (Mosca et al., 2013) (release of February 2014) and iPfam (Mosca et al., 2013) (release of September 2013) pairs of Pfam domains (Finn et al., 2013) that were observed to physically interact in at least one high-resolution 3D co-crystal structure in the Protein Data Bank (Bernstein et al., 1978). Finally, for each PPI we interrogated the Pfam-Pfam interaction data to seek the ppi-mediating domains. When two proteins were experimentally shown to interact and, furthermore, contain interacting Pfam domains, we predict that those Pfam domains are responsible for the interaction and consider them as the interaction interfaces.

Additionally, protein ordered and disordered sequence regions were estimated with the DISO-PRED version2 software using default parameters (Ward et al., 2004). The input protein sequences (fasta files) were downloaded from the UniProt database.

### TCGA and ICGC cancer datasets

Two main data sources were used to retrieve somatic mutations from different cancers: International Cancer Genome Consortium (ICGC, release 15.1) and The Cancer Genome Atlas (TCGA, curated dataset available at Synapse ref. *syn1729383,* (Kandoth et al., 2013)). We discarded those donors with no exonic somatic variants in any protein of the interactome. We discarded also the THCA cancer type from SA due to the abnormal amount of natural germinal variants in the processed file (germinal variants extracted from the 1KP and Spanish MGP individuals) (Consortium et al., 2012; Garcia-Alonso et al., 2014). Finally, donors for whom the number of mutations deviates by three times the standard deviation were excluded.

### Functional characterization of somatic variants

For the merged processed datasets from the ICGC and TCGA/Synapse, we extracted the genome coordinates alongside with the reference and alternative alleles. Next, each variant was mapped to the corresponding transcript/protein and the functional consequence was computed by VARIANT software (Medina et al., 2012). Only point mutations were selected for further analysis.

OncodriveFM (Gonzalez-Perez and Lopez-Bigas, 2012) and OncodriveCLUST (Tamborero et al., 2013) were used to search variants under positive selection across the cohort of tumor samples (pan-can analysis). OncodriveFM was designed to identify genes with a bias towards accumulation of mutations with high functional impact whereas OncodriveCLUST was designed to identify genes with significantly clustered mutations among the gene sequence. IntOGen-mutations pipeline (http://www.intogen.org/mutations/analysis) was used to run OncodriveFM and OncodriveCLUST on the merged mutation datasets (Gonzalez-Perez et al., 2013).

### Statistical analysis of the mutations distribution among protein region types

The distribution of the pancancer nonsynonymous mutations with respect to protein regions was evaluated with a permutation test by comparing the observed mutation count (number of mutations across all donors) in ordered protein sequences, interacting Pfam domains and non-interacting Pfam domains, to a null distribution estimated using a permutation approach. Specifically, the permutation consisted of randomly reassigning mutations to protein sequence positions using all proteins from the structurally resolved interactome, so that the total number of mutations in the interactome is always the same as in the observed case. The p-value can be calculated as the frequency of the observed value in the null distribution.

### Identification of significantly mutated protein interacting interfaces

Identification of PPI interfaces enriched in somatic mutations was assessed by a protein-centric approach. Specifically, statistical significance of mutations in each PPI interface was estimated with one tailed binomial test using overall protein mutation ratio as background. Here, given the observed number of nonsynonymous mutations on a given interacting domain (X), with a given length (L) and a ratio of nonsynonymous mutations along the ordered protein sequence (R,) a one-tailed binomial model was used to compute the probability of observing equal to or greater than *X* mutations in a region L when the null hypothesis is true. We consider that FDR P-values ≤ 0.05 indicate that the ratio of observed mutations in the interacting domain is significantly greater than the ratio of mutation along the ordered protein sequence. Interfaces with less than 5 mutations were discarded.

In order to avoid selecting hyper-mutated sequence regions (Liu et al., 2013), we performed a second test over the synonymous mutations, which are not expected to have a functional implication at the protein level. Interacting interfaces enriched in synonymous mutations were discarded from our predictions since we cannot distinguish whether such bins are just a consequence of hyper-mutation phenomena or, instead, are accumulated due to the conferred selective advantage.

Finally, GOLGA4, RYR2, RYR1 and KRT2 were also excluded from the list as they have been proposed as likely false positives from the methods identifying drivers.

### Survival analysis

Estimation of overall survival for each patient group was calculated using the Kaplan-Meier method implemented in the *survival* package in R. This method uses the clinical variables *donor age at last followup, donor age at diagnosis* and *donor vital status* to quantify the proportion of patients still surviving after a given period of time after its diagnosis. The survival comparison was analyzed using the log-rank test.

### Identification of the minimal connected 3D network mutated in cancer

The interaction network between predicted interaction protein interfaces was created using the SNOW method (Minguez et al., 2009). SNOW detects the largest minimal connected network (MCN) linking all the input proteins and tests if network interconnectivity is significantly greater than the corresponding random expectations. To construct the cancer MCN, we used the structurally resolved interactome allowing the incorporation of one external connecting protein. In order to deal with the structurally resolved version of the interactome, SNOW algorithm was rewritten so that we add only external proteins to participate in the subnetwork if they directly interact with, at least, two enriched interfaces. An empirical distribution of the random expectation of the *average number of nodes per component* parameter for a network of N components was obtained by repeatedly sampling random sets of N protein interfaces from the complete interactome. Then, the real value of the parameter from the MCN between the interfaces of interest is obtained and contrasted with respect to their corresponding random expectations. Finally, the network was visualized using the CellMaps web visualization tool http://cellmaps.babelomics.org/.

## Acknowledgments

Acknowledgements

This work is supported by grants BIO2011-27069 and PRI-PIBIN-2011-1289 from the Spanish Ministry of Economy and Competitiveness (MINECO) and PROMETEO/2010/001 from the Conselleria d’Educaciό of the Valéncia Community. LG-A was supported by fellowship PFIS FI10/00020 from the MINECO. We also thank the support of both initiatives of the ISCIII, MINECO: the National Institute of Bioinformatics (www.inab.org), the CIBER de Enfermedades Raras (CIBERER). We thank the support of Bull through the Bull Chair in Computational Genomics (http://bioinfo.cipf.es/chair_compgenom). We are indebted to the ICGC consortium for making available the data used in this study.

## Author Contributions

LG-A has done the analysis. LG-A and JD have conceived the work and wrote the manuscript.

## Supplementary Information

**Supplementary table 1:** List of predicted cancer interacting interfaces.

## References

Al-Shahrour, F., Díaz-Uriarte, R., and Dopazo, J. (2004). Fatigo: a web tool for finding significant associations of gene ontology terms with groups of genes. Bioinformatics, 20(4):578–580.

Arsenic, R., Lehmann, A., Budczies, J., Koch, I., Prinzler, J., Kleine-Tebbe, A., Schewe, C., Loibl, S., Dietel, M., and Denkert, C. (2014). Analysis of pik3ca mutations in breast cancer subtypes. Applied Immunohistochemistry & Molecular Morphology, 22(1):50–56.

Barbareschi, M., Buttitta, F., Felicioni, L., Cotrupi, S., Barassi, F., Del Grammastro, M., Ferro, A., Dalla Palma, P., Galligioni, E., and Marchetti, A. (2007). Different prognostic roles of mutations in the helical and kinase domains of the pik3ca gene in breast carcinomas. Clinical Cancer Research, 13(20):6064–6069.

Bateman, A., Birney, E., Cerruti, L., Durbin, R., Etwiller, L., Eddy, S. R., Griffiths-Jones, S., Howe, K. L., Marshall, M., and Sonnhammer, E. L. (2002). The pfam protein families database. Nucleic acids research, 30(1):276–280.

Bernstein, F. C., Koetzle, T. F., Williams, G. J., Meyer Jr, E. F., Brice, M. D., Rodgers, J. R., Kennard, O., Shimanouchi, T., and Tasumi, M. (1978). The protein data bank: a computer-based archival file for macromolecular structures. Archives of biochemistry and biophysics, 185(2):584–591.

Brown, C. J., Takayama, S., Campen, A. M., Vise, P., Marshall, T. W., Oldfield, C. J., Williams, C. J., and Keith Dunker, A. (2002). Evolutionary rate heterogeneity in proteins with long disordered regions. Journal of molecular evolution, 55(1):104–110.

Bustin, S. A. (1998). Crosstalk among cancer signalling pathways. Molecular Medicine Today, 4(12):511.

Calleja, V., Laguerre, M., Parker, P. J., and Larijani, B. (2009). Role of a novel ph-kinase domain interface in pkb/akt regulation: structural mechanism for allosteric inhibition. PLoS biology, 7(1):e1000017.

Chen, J. W., Romero, P., Uversky, V. N., and Dunker, A. K. (2006). Conservation of intrinsic disorder in protein domains and families: I. a database of conserved predicted disordered regions. Journal of proteome research, 5(4):879–887.

Chuang, P.-T. and McMahon, A. P. (1999). Vertebrate hedgehog signalling modulated by induction of a hedgehog-binding protein. Nature, 397(6720):617–621.

Consortium,. G. P. et al. (2012). An integrated map of genetic variation from 1,092 human genomes. Nature, 491(7422):56–65.

Consortium, T. U. (2011). Ongoing and future developments at the Universal Protein Resource. Nucleic Acids Research, 39(suppl 1):D214–D219.

Feldman, I., Rzhetsky, A., and Vitkup, D. (2008). Network properties of genes harboring inherited disease mutations. Proceedings of the National Academy of Sciences, 105(11):4323–4328.

Ferlay, J., Steliarova-Foucher, E., Lortet-Tieulent, J., Rosso, S., Coebergh, J., Comber, H., Forman, D., and Bray, F. (2013). Cancer incidence and mortality patterns in europe: estimates for 40 countries in 2012. European journal of cancer, 49(6):1374–1403.

Finn, R. D., Bateman, A., Clements, J., Coggill, P., Eberhardt, R. Y., Eddy, S. R., Heger, A., Hetherington, K., Holm, L., Mistry, J., et al. (2013). Pfam: the protein families database. Nucleic acids research, page gkt1223.

Garcia-Alonso, L., Jiménez-Almazan, J., Carbonell-Caballero, J., Vela-Boza, A., Santoyo-López, J., Antinolo, G., and Dopazo, J. (2014). The role of the interactome in the maintenance of deleterious variability in human populations. Molecular systems biology, 10(9).

Geng, C., He, B., Xu, L., Barbieri, C. E., Eedunuri, V. K., Chew, S. A., Zimmermann, M., Bond, R., Shou, J., Li, C., et al. (2013). Prostate cancer-associated mutations in speckle-type poz protein (spop) regulate steroid receptor coactivator 3 protein turnover. Proceedings of the National Academy of Sciences, 110(17):6997–7002.

Gerber, A., Wilson, C., Li, Y., and Chuang, P. (2006). The hedgehog regulated oncogenes gli1 and gli2 block myoblast differentiation by inhibiting myod-mediated transcriptional activation. Oncogene, 26(8):1122–1136.

Goh, K., Cusick, M., Valle, D., Childs, B., Vidal, M., and Barabèsi, A. (2007). The human disease network. Proceedings of the National Academy of Sciences, 104(21):8685–8690.

Gonzalez-Perez, A. and Lopez-Bigas, N. (2012). Functional impact bias reveals cancer drivers. Nucleic acids research, page gks743.

Gonzalez-Perez, A., Perez-Llamas, C., Deu-Pons, J., Tamborero, D., Schroeder, M. P., Jene-Sanz, A., Santos, A., and Lopez-Bigas, N. (2013). Intogen-mutations identifies cancer drivers across tumor types. Nature methods.

Hanahan, D. (2000). The hallmarks of cancer. Cell, 100(1):57–70.

Hanahan, D. and Weinberg, R. A. (2011). Hallmarks of cancer: the next generation. Cell, 144(5):646–674.

Javelaud, D., Alexaki, V.-I., Pierrat, M.-J., Hoek, K. S., Dennler, S., Van Kempen, L., Bertolotto, C., Ballotti, R., Saule, S., Delmas, V., et al. (2011). Gli2 and m-mitf transcription factors control exclusive gene expression programs and inversely regulate invasion in human melanoma cells. Pigment cell & melanoma research, 24(5):932–943.

Jonsson, P. F. and Bates, P. A. (2006). Global topological features of cancer proteins in the human interactome. Bioinformatics, 22(18):2291–2297.

Kalinsky, K., Jacks, L. M., Heguy, A., Patil, S., Drobnjak, M., Bhanot, U. K., Hedvat, C. V., Traina, T. A., Solit, D., Gerald, W., et al. (2009). Pik3ca mutation associates with improved outcome in breast cancer. Clinical Cancer Research, 15(16):5049–5059.

Kandoth, C., McLellan, M. D., Vandin, F., Ye, K., Niu, B., Lu, C., Xie, M., Zhang, Q., McMichael, J. F., Wyczalkowski, M. A., et al. (2013). Mutational landscape and significance across 12 major cancer types. Nature, 502(7471):333–339.

Lai, Y.-L., Mau, B.-L., Cheng, W.-H., Chen, H.-M., Chiu, H.-H., and Tzen, C.-Y. (2008). Pik3ca exon 20 mutation is independently associated with a poor prognosis in breast cancer patients. Annals of surgical oncology, 15(4):1064–1069.

Li, C., Ao, J., Fu, J., Lee, D.-F., Xu, J., Lonard, D., and O’Malley, B. W. (2011). Tumor-suppressor role for the spop ubiquitin ligase in signal-dependent proteolysis of the oncogenic co-activator src-3/aib1. Oncogene, 30(42):4350–4364.

Liu, L., De, S., and Michor, F. (2013). Dna replication timing and higher-order nuclear organization determine single-nucleotide substitution patterns in cancer genomes. Nature communications, 4:1502.

Mangone, F. R., Bobrovnitchaia, I. G., Salaorni, S., Manuli, E., and Nagai, M. A. (2012). Pik3ca exon 20 mutations are associated with poor prognosis in breast cancer patients. Clinics, 67(11):1285–1290.

McLendon, R., Friedman, A., Bigner, D., Van Meir, E. G., Brat, D. J., Mastrogianakis, G. M., Olson, J. J., Mikkelsen, T., Lehman, N., Aldape, K., et al. (2008). Comprehensive genomic characterization defines human glioblastoma genes and core pathways. Nature, 455(7216):1061–1068.

Medina, I., Carbonell, J., Pulido, L., Madeira, S., Goetz, S., Conesa, A., Tárraga, J., Pascual-Montano, A., Nogales-Cadenas, R., Santoyo, J., et al. (2010). Babelomics: an integrative platform for the analysis of transcriptomics, proteomics and genomic data with advanced functional profiling. Nucleic acids research, 38(suppl 2):W210.

Medina, I., De Maria, A., Bleda, M., Salavert, F., Alonso, R., Gonzalez, C. Y., and Dopazo, J. (2012). Variant: Command line, web service and web interface for fast and accurate functional characterization of variants found by next-generation sequencing. Nucleic acids research, 40(W1):W54–W58.

Minguez, P., Gotz, S., Montaner, D., Al-Shahrour, F., and Dopazo, J. (2009). Snow, a web-based tool for the statistical analysis of protein-protein interaction networks. Nucleic Acids Research, 37(suppl 2):W109.

Mosca, R., Céol, A., Stein, A., Olivella, R., and Aloy, P. (2013). 3did: a catalog of domain-based interactions of known three-dimensional structure. Nucleic acids research, page gkt887.

Oro, A. E., Higgins, K. M., Hu, Z., Bonifas, J. M., Epstein, E. H., and Scott, M. P. (1997). Basal cell carcinomas in mice overexpressing sonic hedgehog. Science, 276(5313):817–821.

Parikh, C., Janakiraman, V., Wu, W.-I., Foo, C. K., Kljavin, N. M., Chaudhuri, S., Stawiski, E., Lee, B., Lin, J., Li, H., et al. (2012). Disruption of ph-kinase domain interactions leads to oncogenic activation of akt in human cancers. Proceedings of the National Academy of Sciences, 109(47):19368–19373.

Porta-Pardo, E. and Godzik, A. (2014). e-driver: a novel method to identify protein regions driving cancer. Bioinformatics, page btu499.

Reimand, J. and Bader, G. D. (2013). Systematic analysis of somatic mutations in phosphorylation signaling predicts novel cancer drivers. Molecular systems biology, 9(1).

Richter, J., Schlesner, M., Hoffmann, S., Kreuz, M., Leich, E., Burkhardt, B., Rosolowski, M., Ammerpohl, O., Wagener, R., Bernhart, S. H., et al. (2012). Recurrent mutation of the id3 gene in burkitt lymphoma identified by integrated genome, exome and transcriptome sequencing. Nature genetics, 44(12):1316–1320.

Ruch, J. M. and Kim, E. J. (2013). Hedgehog signaling pathway and cancer therapeutics: progress to date. Drugs, 73(7):613–623.

Schmitz, R., Young, R. M., Ceribelli, M., Jhavar, S., Xiao, W., Zhang, M., Wright, G., Shaffer, A. L., Hodson, D. J., Buras, E., et al. (2012). Burkitt lymphoma pathogenesis and therapeutic targets from structural and functional genomics. Nature, 490(7418):116–120.

Tamborero, D., Gonzalez-Perez, A., and Lopez-Bigas, N. (2013). Oncodriveclust: exploiting the positional clustering of somatic mutations to identify cancer genes. Bioinformatics, 29(18):2238–2244.

Vogelstein, B., Papadopoulos, N., Velculescu, V. E., Zhou, S., Diaz, L. A., and Kinzler, K. W. (2013). Cancer genome landscapes. science, 339(6127):1546–1558.

Wang, X., Wei, X., Thijssen, B., Das, J., Lipkin, S. M., and Yu, H. (2012). Three-dimensional reconstruction of protein networks provides insight into human genetic disease. Nature biotechnology, 30(2):159–164.

Ward, J. J., McGuffin, L. J., Bryson, K., Buxton, B. F., and Jones, D. T. (2004). The disopred server for the prediction of protein disorder. Bioinformatics, 20(13):2138–2139.

Wong, J. M., Ionescu, D., and Ingles, C. J. (2003). Interaction between brca2 and replication protein a is compromised by a cancer-predisposing mutation in brca2. Oncogene, 22(1):28–33.

Wu, W.-I., Voegtli, W. C., Sturgis, H. L., Dizon, F. P., Vigers, G. P., and Brandhuber, B. J. (2010). Crystal structure of human akt1 with an allosteric inhibitor reveals a new mode of kinase inhibition. PLoS One, 5(9):e12913.

Yu, H., Braun, P., Yildirim, M. A., Lemmens, I., Venkatesan, K., Sahalie, J., Hirozane-Kishikawa, T., Gebreab, F., Li, N., Simonis, N., et al. (2008). High-quality binary protein interaction map of the yeast interactome network. Science, 322(5898):104–110.

Zhang, L., Shi, L., Zhao, X., Wang, Y., and Yue, W. (2013). Pik3ca gene mutation associated with poor prognosis of lung adenocarcinoma. OncoTargets and therapy, 6:497.

